# Structural interactions between pandemic SARS-CoV-2 spike glycoprotein and human Furin protease

**DOI:** 10.1101/2020.04.10.036533

**Authors:** Naveen Vankadari

## Abstract

The SARS-CoV-2 pandemic is an urgent global public health emergency and warrants investigating molecular and structural studies addressing the dynamics of viral proteins involved in host cell adhesion. The recent comparative genomic studies highlight the insertion of Furin protease site in the SARS-CoV-2 spike glycoprotein alerting possible modification in the viral spike protein and its eventual entry to host cell and presence of Furin site implicated to virulence. Here we structurally show how Furin interacts with the SARS-CoV-2 spike glycoprotein homotrimer at S1/S2 region, which underlined the mechanism and mode of action, which is a key for host cell entry. Unravelling the structural features of biding site opens the arena in rising bonafide antibodies targeting to block the Furin cleavage and have great implications in the development of Furin inhibitors or therapeutics.

## Introduction

The pandemic Corona Virus Disease 2019 (COVID-19) caused by Severe Acute Respiratory Syndrome Coronavirus 2 (SARS-CoV-2), is an urgent public health emergency and made a serious impact on global health and economy (1). To date, more than 86,000 deaths and 1.5 million confirmed positive cases were reported globally, making the most contagious pandemic in the last decade (www.coronavirus.gov). Since the initial reports on this pneumonia-causing novel coronavirus (SARS-CoV-2) in Wuhan, China, mortality and morbidity are increasing exponentially around the globe despite several antiviral and antibody treatments (2). Most available neutralising antibodies in use are targeting the SARS-CoV-2 spike glycoprotein, which is essential for host cell adhesion via ACE2 and CD26 receptors (3, 4), but infection control is still insignificant. Meanwhile, several antiviral drugs (Ritonavir, Lopinavir, Chloroquine, Remdesivir and others) targeting different host and viral proteins are been clinically evaluating and repurposing to combat SARS-CoV-2 infection (2, 5). With the drastic increasing number of the positive cases around the world (www.cdc.gov), moderate response to antivirals under clinical trials and poor response to antibodies targeting spike SARS-CoV-2 spike glycoprotein is a serious concern and warrants detail understanding of the molecular and structural features of SARS-CoV-2 structural proteins in native condition and post-viral infection. This will abet in understanding the dynamics and mechanism of viral action on the human cell.

In this regard, several epidemiological and evolutionary reports have highlighted the several unique sequence deletions and insertions in the SARS-CoV-2 genome compare to previous known SARS and Bat coronavirus (6, 7). Among the various genetic variations, insertion of Furin protease cleavage site in the spike glycoprotein (aa682 – aa689) is strikingly novel in SARS-CoV-2 (4, 8) and has not been found in other related coronaviruses (SARS-CoV-1, Bat-CoV, Pangolin) (MERS contain pseudo binding site) (Fig. S1A). Furin protease belongs to the family of calcium (Ca2+)-dependent proprotein/prohormone convertase (PCs) which is ubiquitously expressed in humans but its levels are elevated lung cystic fibrosis (9). Furin protease also cycles from trans-Golgi network (TGN) to cell membrane (virus attaches) and endosomes (virus translocate in endosomes). Interestingly, Furin and other related proteases are highly specific and known for cleaving different viral (Influenza, HIV) envelope glycoproteins, thereby enhancing viral fusion with the host cell membrane (10). Furthermore, Furin preferentially recognizes the motif R-X-K/R-R and cleaves the peptide in the presence of Ca2+, which is physiologically connected to different viral infections (10, 11). However, about SARS-CoV-2, it is elusive that how Furin could bind and act on the viral spike glycoprotein. Hence to understand the interaction mode and mechanism of Furin action over the spike glycoprotein warrants further structural and biomolecular studies.

## Methods

Considering the current public health crisis and to better understand the structural and molecular mode of interactions between SARS-CoV-2 spike protein and human Furin, we resolve the structure of SARS-CoV-2 spike glycoprotein in complex with Furin protease via molecular dynamics and simulations. Unfortunately, the only two available SARS-CoV-2 spike glycoprotein Cryo-EM structures (PDB: 6VSB and 6VXX) are incomplete and has several gaps in the built structure and also lacks the structure for Furin cleavage sites (3, 12). As these EM structures built on molecular replacement with SARS-CoV-1 (PDB: 6ACG), Furin cleavage sites in the spike protein is flexible and novel insertion only in the SARS-CoV-2, the EM structures lack this important region. Hence, for the molecular dynamics and simulation studies, we directed to use previously published and validated model structure of full-length SARS-CoV-2 spike glycoprotein (4) and published structure of human Furin (PDB: 1P8J or 1JXH) (11). The RMSD of the previously published model structure and Cryo-EM structure was 0.84, which suggests overall structural accuracy even with the presence of Furin cleavage sites. The binding free energies were taken into consideration for selecting the best possible model. Further validation and refinement was completed by ensuring that the residues occupied Ramachandran favoured positions using Coot (www.mrc-imb.cam.uk/). The final complex structure was then compared with the initial Furin structure and their overall RMSD was found to be 0.28 Å for Ca atoms.

## Results

The overall complex structure shows three Furin proteases binding to the mid or equatorial region (mid region of S1 and S2 domain (S1/S2)) of SARS-CoV-2 spike glycoprotein homo-trimer at the off-centric and adjacent side of spike trimer (Fig. 1 and S1B). The binding Furin proteases adopt a clamp-like fashion, where it clips to the cleavage site of the spike glycoprotein. Furthermore, the binding of Furin protease creates a large burred interface of ~1,100Å2 (~368Å2/Furin) between the proteins, as calculated from the PISA server (https://www.ebi.ac.uk/pdbe/pisa/). This suggests a bonafide and tight interaction of Furin protease over the spike glycoprotein and Furin. The depth, shape and charge of Furin protease are well known and it has canyon-like crevice and its active site pocket is conserved in many species and the catalytic or substrate-binding pocket is made of key amino acid residues R185, M189, D191, N192, R193, E229, V231, D233, D259, K261, R298, W328 and Q346 (10, 11) (Fig. 2 and S2). Interestingly, these residues are also well-positioned to interact with the viral spike protein cleavage site in our complex structure and the entire substrate-binding pocket of Furin protease appears like a canyon-like crevice, which can accommodate a large portion of target protein/peptide. The results show that the SARS-CoV-2 spike glycoprotein amino acid residues N657 to Q690 are the prime interacting residues with the Furin protease. The position and orientation of these unique residues involved in Furin recognition are well exposed and organise in a flexible loop. The spike protein residues N657, N658, E661, Y660, T678, N679, S680, R682, R683, R685, S689, Q690 makes the strong interaction with the Furin protease (Fig. 2A). The interaction between the viral spike glycoprotein and Furin protease is mediated via several van der Waals or by hydrogen bonding. Furthermore, the entire cleavage loop of viral spike protein fits into the canyon-like substrate-binding pocket of Furin protease. It is quite interesting to notice that none of the previously known coronaviruses had this novel Furin protease cleavage site in the spike glycoprotein, which accentuates the novelty and uniqueness of SARS-CoV-2. In addition, previous reports on the glycosylation of spike glycoprotein show that Furin cleavage site in the SARS-CoV-2 spike glycoprotein is not targeted by the glycosylation, hence this cleavage loop is completely solvent-exposed (4). This further corroborates the potential attack of Furin protease over the S1/S2 cleavage site in the SARS-CoV-2 spike glycoprotein. Based on the Furin binding mode and structural interaction, we propose the following supposition. The binding and cleaving (priming) the spike glycoprotein at S1/S2 region by Furin protease might cut the spike glycoprotein into N-terminal S1 domain involved in host cell recognition and C-terminal S2 membrane-anchored domain involved in host cell penetration and entry, thus making the SARS-CoV-2 highly virulent. In support of this supposition, it is evident in infectious bronchitis virus that presence of Furin cleavage site has pronounced virulence suggesting Furin cleavage increase the virulence (13).

**Figure 1:**
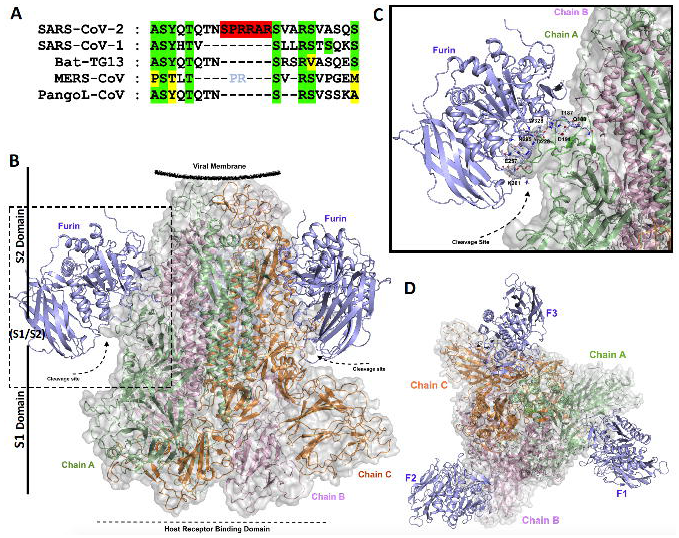
**(A)** Overall structure showing SARS-CoV-2 spike glycoprotein homo-trimer (substrate un-bound or closed conformation) in complex with human Furin protease. The three monomers of SARS-CoV-2 spike glycoprotein homo-trimer are coloured in green (Chain A), pink (Chain B) and orange (Chain C) and the Furin protease is coloured in blue. The spike protein cleavage site is indicated by arrow and S1/S2 domain are labelled accordingly. **(B)** Enlarged view showing the single Furin interacting with its target cleavage site (loop) of SARS-CoV-2 spike glycoprotein. Colour coding and labelling is same as above. **(C)** Top view of Figure 1A, showing the SARS-CoV-2 spike glycoprotein homotrimer bound to three Furin proteases at the adjoining conformation at the S1/S2 region.

**Figure 2:**
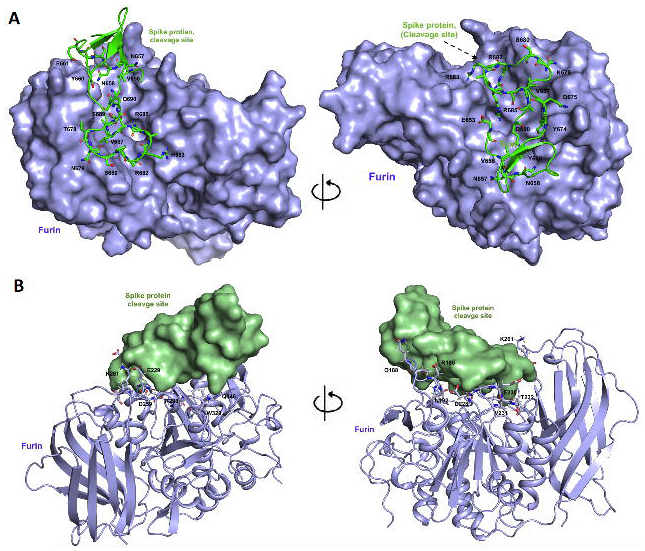
Surface and cartoon model showing the detailed amino acid interaction between the Furin protease and SARS-CoV-2 spike glycoprotein. **(A)** Front and orthogonal view of Furin (blue, surface) interacting with the SARS-CoV-2 spike glycoprotein (green, sticks). For clear visualization one the Furin binding loop is shown. The canyon-like crevice is distinguishable in Furin and the side chin residues of spike protein are labelled accordingly. **(B)** Front and orthogonal view of Furin (blue, sticks and cartoon) interacting with target S1/S2 cleavage site of SARS-CoV-2 spike glycoprotein (green, surface). The key residues of Furin involved in the interaction with S1/S2 cleavage site are shown in sticks and labelled accordingly.

## Discussion

Based on this enzyme cleavage action and separation of N- and C-terminal domain of spike glycoproteins also could make the ACE2 and CD26 inhibitors of least effective, as upon cleavage the N-terminal S1 domains are not required for the cell penetration. This also raises a caution that while making neutralizing antibodies targeting SARS-CoV-2 spike glycoprotein, these cleavage activities need to be considered. Hence, we speculate that antibodies against S2 domain and drugs targeting S1 trimerization could be more promising. These observations and structure-guided molecular interaction with novel Furin protease guide us to suggest that SARS-CoV-2 have different infection modes with that of earlier known coronaviruses. Repurposing and developing targets (inhibitors and peptide) to block the Furin protease found to be another potential therapeutic option and also warrant clinical investigation. This study also first to show structurally that how the human Furin interacts with the coronavirus spike glycoprotein, which underlines its mechanism of action. This structural and molecular dynamics study has great implications to further develop Furin protease inhibitors to block the protease activity of Furin and also abet in the development of bonafide antibodies targeting the S1/S2 Furin cleavage site of spike glycoprotein and accenture the development of future therapeutics.

## Acknowledgements

I thank the Monash University Software Platform for licence access to the concerned software. I also acknowledge Joseph Polidano of University of Melbourne for editing and proof reading the manuscript.

